# Raman flow cytometry using time delay integration

**DOI:** 10.1101/2024.10.17.617595

**Authors:** Matthew Lindley, Toshiki Kubo, Stéphanie Devineau, Menglu Li, Jing Qiao, Takuya Yashiro, Shiroh Iwanaga, Kazuyo Moro, Katsumasa Fujita

**Author notes:** These authors contributed equally to this work (Matthew Lindley, Toshiki Kubo). Dartmouth College, Thayer School of Engineering, 15 Thayer Drive, Hanover, NH 03755, USA. Shenzhen Medical Academy of Research and Translation, Shenzhen 518107, China (Menglu Li).

## Abstract

Raman flow cytometry offers chemically sensitive, label-free measurement of cells and particles; however, the technique suffers from low cell throughput due to the weak Raman signal. Here, we demonstrate the use of time-delay integration to achieve Raman flow cytometry combined with dual-sided line illumination. The use of line illumination from both sides of the cell flow capillary kept the cell stream in the detection area by balancing optical force from the illumination lines. The time-delay integration allowed accumulation of Raman signals from flowing cells without sacrificing the spectrum readout rate. With the developed system, we achieved Raman flow cytometry at throughputs of 32 and 75 events per second for cell and particle detection, respectively. We applied the technique for analyzing lipid uptake in HepG2 cells and degranulation in bone marrow-derived murine mast cells.

## Introduction

Chemically sensitive flow cytometry enables rapid identification and quantification of cellular phenotype for large cell samples at single cell resolution. It is used widely in biological, medical and industrial fields, where applications include cell phenotyping, disease discovery, and rare cell detection.^1^ Commonly in these applications, flow cytometry distinguishes phenotype by detecting exogenous fluorophore “labels” that are bound to target content within the cell or by associating transgenic fluorescent protein expression with target cellular processes. While this has enabled numerous advancements in understanding cell biology, such methods can suffer from non-specific labeling, cell toxicity, and incompatibility with therapeutic applications. These challenges have given rise to “label-free” flow cytometry, which identifies target chemical content in the cell via its direct interaction with light.

Raman flow cytometry seeks to realize label-free measurement of intracellular content by detecting the inelastic scattering of light from molecular vibration. In this process, the scattered light is given an energy-shift, referred to as “Raman shift,” that is conserved by the gain or loss of molecular vibration in the sample. The energies of these vibrational modes are determined by molecular structure, which allows Raman spectroscopy to produce a spectrum that serves as a “fingerprint’ for identifying molecules in a specimen. Raman flow cytometers have been used to detect phenotype based on intracellular carotenoid^2^, polysaccharide^3,4^, and lipids and protein^5,6^ content, as well as to analyze cellular metabolism^2^, differentiation^5^, size^7^, and morphology^8^. Additionally, several Raman flow cytometers have been equipped with real-time analysis and mechanical response to allow Raman-activated cell sorting based on enzyme function^9,10^, intracellular chemical content^11,12^, and morphology^6^.

For Raman flow cytometers one of two kinds of Raman scattering spectroscopy are employed: spontaneous or coherent. Spontaneous Raman scattering spectroscopes are typically less expensive and simpler to build. However, they produce weak signal due to the low cross section of Raman scattering, which is roughly fifteen orders of magnitude smaller than that of fluorescence^13^. Because of this, measurement usually requires a few hundred milliseconds of acquisition for strongly scattering targets up to several seconds of acquisition for weakly scattering targets. To facilitate such long acquisition times, spontaneous Raman flow cytometers and flow-based sorters have relied on trapping cells during measurement^9–11,14–17^. This adds significant complexity to the cytometers and reduces throughput. To the best of our knowledge, spontaneous Raman flow cytometers demonstrated to date are limited to throughputs of less than 5 events/s^17^, where an event is defined as the detection and measurement of a cell, cell cluster, or cell-sized debris. Raman scattering signal can be enhanced by using resonance Raman measurement, which relies on strong optical absorption by the analyte to increase scattering probability, but this limits addressable analytes to strongly absorbing intracellular chemicals such as carotenoids and chlorophyll^18^. Thus, low throughput has remained a bottleneck for spontaneous Raman flow cytometers, preventing applications requiring rapid analysis or rare cell detection. A second type of Raman scattering spectroscopy, called coherent Raman scattering spectroscopy, makes use of pulsed lasers to drive a vibrational coherence in the sample during measurement. For short acquisitions, this produces much better signal to noise ratios than spontaneous Raman measurement, though at typically increased cost and experimental complexity. The use of coherent Raman scattering has allowed throughputs up to ∼2000 event/s in Raman flow cytometers^2^, however, coherent Raman flow cytometers to date are limited by a smaller spectral bandwidth compared to their spontaneous Raman counterparts, which reduces the number of chemical species they can detect.

Here we demonstrate the use of time-delay-integration (TDI) to overcome the throughput bottleneck of spontaneous Raman-based flow cytometry while maintaining its broadband spectral content. TDI is a technique developed to counteract motion blurring of an image as it transits the face of a charge-coupled device (CCD) array during signal acquisition. It accomplishes this by shifting the photoelectrons integrated during a continuous acquisition along the CCD array at a rate that matches the motion of the sample’s image (Figure 1A), with the caveat that the velocity of the image is uniform and unidirectional. TDI was first demonstrated in astronomy^19^, with subsequent use in radiographic and CT imaging^20^, capillary electrophoresis^21^, and fluorescence flow cytometry^22^. By applying TDI to Raman flow cytometry, we demonstrate a trapping-free spontaneous Raman flow cytometer capable of cell measurement of various cell types at up to 32 events/s and particle measurement at 75 events/s with a spectral range from spanning the fingerprint, silent, and CH-stretch Raman regions (500 – 3300 cm^-1^) with a spectral resolution of <5 cm^-1^. We typify our flow cytometer’s performance with the measurement of microplastic particles and different cell types, namely HeLa cells, hepatocytes, mast cells, and red blood cells. Then we demonstrate analysis of lipid accumulation in HepG2 cells grown under different culture conditions. Finally, we discuss the theoretical limits to throughput with this regime.

**Figure 1:**
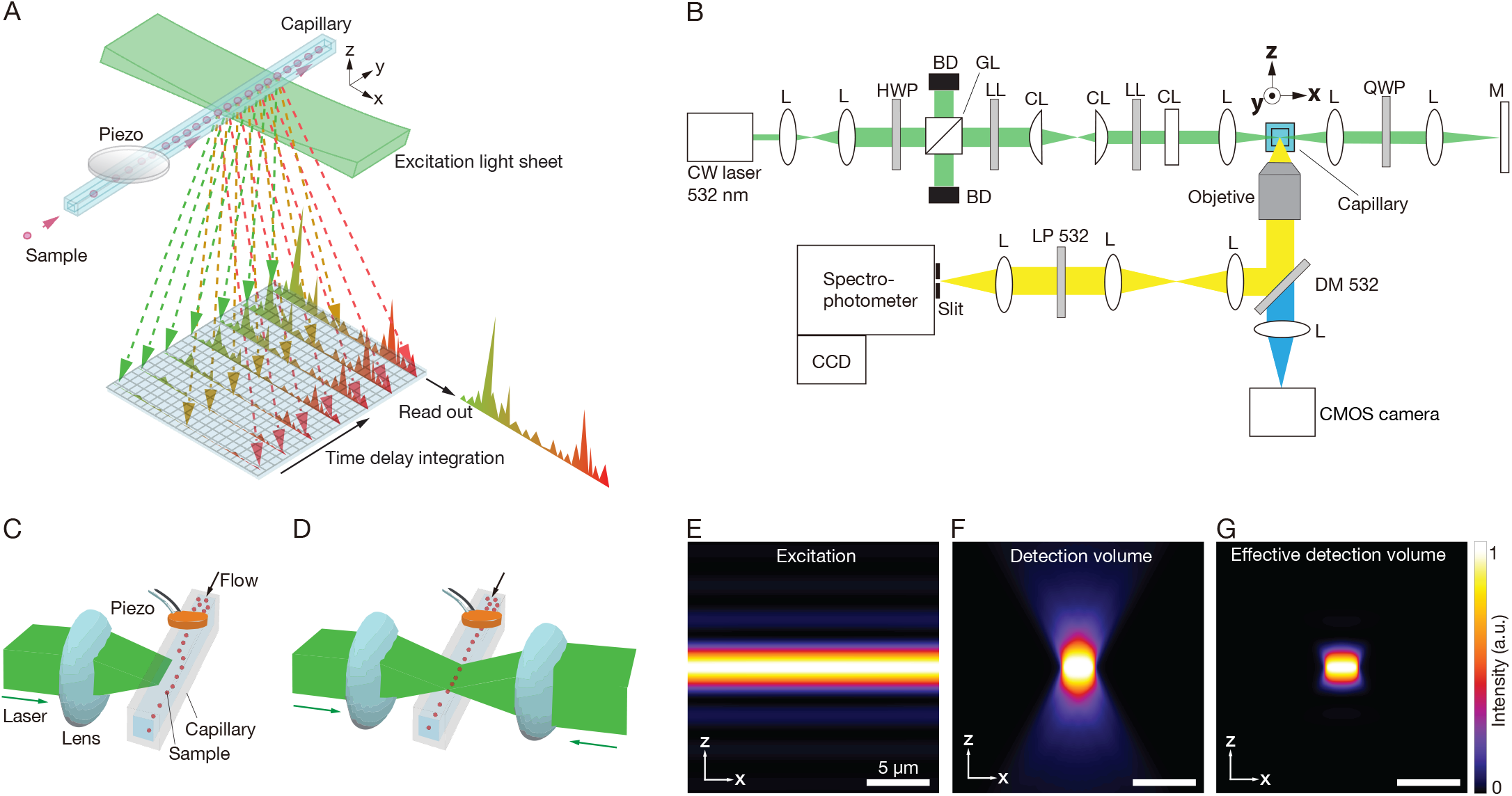
Diagram of TDI Raman flow cytometer & optics simulations. **(A)** TDI Raman flow cytometry scheme. **(B)** Optical layout; L, CL, HWP, GL, BD, LL, QQP, M, DM, SP, and LP are lens, cylindrical lens, half wave plate, Glan-laser Calcite Polarizer, beam dump, laser line filter, quarter wave plate, mirror, dichroic mirror, short-pass edge filter, and long-pass edge filter, respectively. The direction of flow is along the y-direction. (C-D) Illustration of flow path with **(C)** one- and **(D)** two-way illumination. The cell flow is focused to the capillary center using a piezo transducer. When having one-way illumination, the laser pushes cells out of alignment with the measurement region. Applying counter-propagating illumination enables balancing the pressure on the cells. **(E to G)** Simulated distribution of **(E)** intensity of excitation light sheet, **(F)** detection volume given by the convolution product of the detection PSF and confocal slit, and **(G)** effective detection volume considering both excitation and detection volumes. We assumed a slit width of 50 µm, which corresponds to 5.2 µm at the object space.

## Results

### Flow cytometer design

Our flow cytometer consisted of three parts: the optical illumination line, the optical detection line, and the flow path, as shown in Figure 1B. The optical illumination line contained a high power 532 nm continuous wave (CW) laser with optics that shaped the beam into a light sheet, aligned such that the long axis of its focal waist was colinear with the flow path. We calculated light sheet intensity in the focal region to range from 0.26 – 3.2 mW/µm^2^ depending laser output settings, based on a simulation of beam geometry and power measurements at the sample position. The light sheet imparted a force upon flowing particles and cells in the direction of optical propagation, which tended to push the sample out of alignment with the measurement region. To counteract this, we retroreflected the illumination line as shown in Figure 1C & D, such that the flowing sample was illuminated (pushed) from both sides.

We collected scattering signal using an objective lens mounted orthogonally to the light sheet’s propagation and long focal waist axes. We used a tube lens, dichroic mirror, and relay lenses to carry the Raman scattering signal from the objective to a spectrometer equipped with a cooled CCD camera oriented so that the images of flowing cells travelled along the spectrometer slit. We then oriented the spectrometer’s CCD array to align its TDI axis (400 pixels) colinearly with motion in the imaging plane. We used the CCD array’s orthogonal axis (1340 pixels) as the Raman shift axis following dispersion by the spectrometer’s grating.

The measurement region in our flow cytometer can be modeled as a product of the point-spread functions (PSFs) of our illumination and detection optics. We designed the image magnification at the CCD camera to be low (4.8 to 9.6 times) so that signal from a relatively large volume within the sample could be integrated in a single CCD pixel, with the goal of minimizing readout noise per signal. Figure 1E shows a model of the intensity distribution of the excitation light sheet at the measurement position. A model of our detection PSF, based on system optics, slit width, and CCD pixel size is shown in Figure 1F. The product of these two volumes gives the effective measurement region, shown in Figure 1G. For TDI measurement of Raman scattering from the cells, it was imperative that the cells travelled along the measurement region. We flowed cells through a square quartz capillary (outer width and height 600 µm, channel width and height 240 µm) using acoustic focusing from an attached piezoelectric transducer to confine cell motion to the center of the capillary (Figure 1B) and then adjusted the capillary position relative to the light sheet to align cell motion with the measurement region.

### Theoretical treatment of throughput

A brief theoretical treatment of throughput follows. Acquisition time τ is given by the time it takes a cell image to transit the length of the CCD array, such that 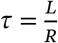, where *L* is pixel number along the TDI axis and *R* is the spectral rate (here we assume a well-controlled flow speed matches image transit across the CCD face to the spectral rate). Throughput *T* maximizes as 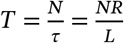, where *N* is the expectation value of the number of cells that can be distributed along the CCD’s TDI axis. This number is determined by sample concentration, which increases as magnification is reduced. Ideally, at low magnification the entire signal from a cell is confined to just one pixel-width along the direction of TDI transit. Distributing signal among a minimum number of pixels also carries the benefit of minimizing the CCD readout noise per event. In an optimal case, single cells could be distinguished by 2 pixels in a cell-blank-cell-blank pattern along the TDI-axis, where 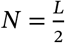. In such a case, maximum throughput becomes dependent only on spectral rate, 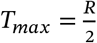, provided the CCD array is long enough for acceptable signal accumulation and the cell distribution is well controlled. However, for the simple flow channel and acoustic focusing design used in this experiment the spatial distribution of cells along the flow axis was not controlled. Under such cases, intercell spacing is commonly modelled using a Poisson distribution, 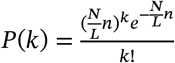, where *P*(*k*) is the probability that *k* cells are observed in a pixel interval *n*,, provided there is no cell-to-cell influence. In this regime single cell event throughput is 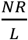 and *P*(1) maximizes at 37%.

### Microplastic particle measurements

To typify flow cytometer performance, we performed measurement of 3 µm polystyrene beads by our flow cytometer and achieved a sustained throughput of 75 events/s (Movie S1). The bead sample was prepared at 6×10^6^ particles/mL, though analysis of the highspeed video shows an average of 116 particles/s, equivalent to 3.3×10^6^ particles/mL, transiting the capillary. We attribute the reduction in number density between sample preparation and measurement to beads settling out of suspension in the pump reservoir during the setup and execution of the flow cytometry.

The difference in particle count between our high-speed video and our spectrometer can be explained by comparing their temporally aligned data. An image composited from three frames of high-speed video of beads undergoing measurement is shown in Figure 2A. In the image the beads flow from right to left and the direction of light sheet propagation is from top to bottom for the initial illumination. The beads scattered a higher portion of this light than the flow media, causing them to appear as white particles against a dark background in the image. The spectral readout of Raman scattering from the beads as captured by our TDI spectrometer is shown in Figure 2B, following background removal. Here an increase in baseline intensity indicates a particle transit event. The spectral data in Figure 2B was captured simultaneously with the video data in 2A, and the data have been arranged to temporally align in the figure. Pink arrows indicate beads flowing far enough apart that their spectra could be easily resolved as separate events. The blue arrow indicates two beads clumped together such that their spectra begin to overlap temporally. Additionally, imperfect acoustic focusing allowed some beads to transit the measurement region at the edge of the light sheet where they were largely outside the collection optics PSF. Scattering from these out-of-focus beads (an example is indicated by the green arrow) appears in the high-speed camera images but was largely rejected by the slit of our spectrometer, which provided a high sectioning capability. The characteristic fingerprint-region peaks of polystyrene around 1000 cm^-1^, 1600 cm^-1^, and the CH-stretch region peaks on either side of 3000 cm^-1^ are readily apparent in the spectra of in-focus beads but missing for out-of-focus beads.

**Figure 2:**
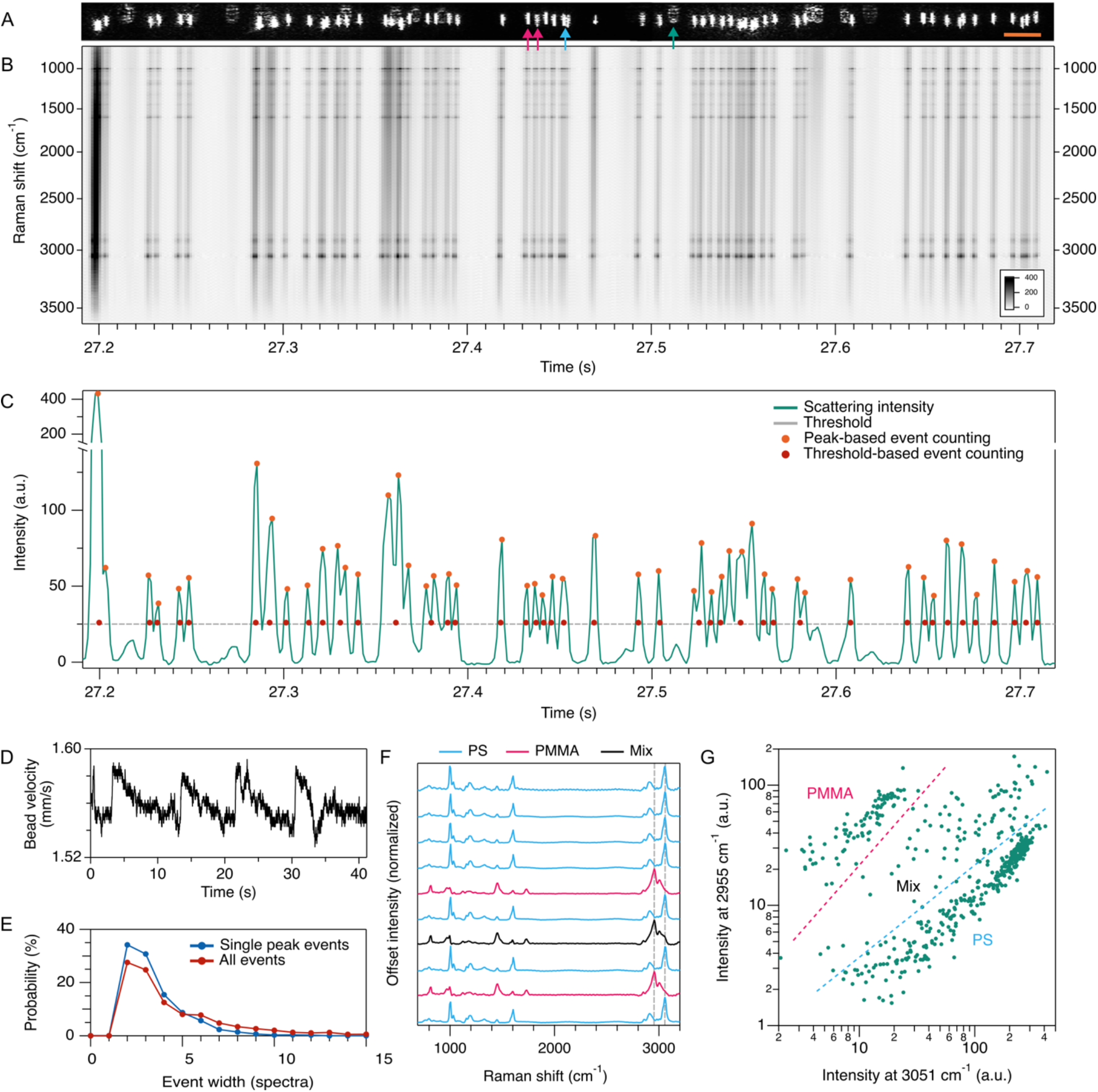
High throughput measurement of PS microbeads and measurement of a mixed PS-PMMA beads sample. **(A)** Images of 3 µm polystyrene beads flowing in capillary at ∼140 beads per second, captured by high-speed camera. Scale bar 30 µm. Pink and blue arrows denote successive beads which can be distinguished or not be distinguished as individual events, respectively, in the spectral readout. Green arrow indicates a bead flowing outside the optical focal plane. **(B)** Corresponding spectral readout from flow cytometer after denoising. **(C)** Integrated scattering signal, with events selected by a peak-based algorithm (orange circles) or a threshold-based algorithm (red circles). **(D)** Spectral width of events at the threshold for all events (N=2271). (E) Histogram of event width at threshold indicated in C. **(F)** Successive single event spectra from a mixed PS and PMMA sample. Vertical gray lines show peak positions plotted in G. **(G)** Scatter plot of Raman shift intensity at the primary PS and PMMA peaks in the CH stretch region.

We tested two algorithms for detecting and counting events in the spectral data, detailed in Figure 2C, which shows the integrated signal intensity output by the spectrometer for the temporal region corresponding to 2A & B in green. The first event selection algorithm defined events by a simple intensity threshold, where single events were denoted by signal rising above then falling below an arbitrarily chosen threshold value, shown by the grey line in Figure 2C. Events selected by this algorithm are displayed by a blue dot at the temporal center of the event, and we produced single event by averaging above-threshold spectra for each single event. This algorithm often combined spectra from multiple beads into a single event, particularly for a low threshold value, so we developed a more robust peak-based event selection algorithm. Our peak-based algorithm selected peak positions in the temporal intensity data based on zero-crossing of the derivative, with below-threshold peaks discarded. Single event spectra were then averaged from the set of 3 to 11 spectra (as set by the user) spanning each peak position. Example peak selections by this algorithm are shown by the orange dots in Figure 2C.

The data chosen for Figure 2A, B, and C consist of a temporal region of data of data with many beads in immediate proximity to one another, making the data useful for judging TDI performance as throughput is maximized. The composite image shows 70 beads transiting the measurement region in 0.5 s, equivalent to 140 beads/s. Of these, 12 beads (17%) are out of focus, leaving 58 (116 beads/s) capable of being spectrally detected. The temporally aligned TDI spectral readouts in Figure 2B & C show our threshold-based algorithm detecting 43 events (86 events/s) and our peak-based algorithm detecting 51 events (102 events/s). For the latter, 7 events are “multi-bead” events in which the spectra from two closely spaced beads are indistinguishable, giving a yield of 86% for single bead events. The data shown in Figure 2A, B, & C were selected as the extreme case from a dataset of 41 s of continuous measurement, across which threshold-based selection reported a throughput of 55 events/s and peak-based selection returned 75 events/s. These results from peak-based selection are quite close to the theoretical treatment of throughput at this spectral rate and bead number density for a pixel interval of 3, which matches bead diameter. Under these parameters, theory suggests a throughput of 72 events/s, with a single bead yield of 37% of spectra and 78% of events. The higher rate of single bead event yield in our empirical data suggests that bead-to-bead interactions pushes us slightly out of the Poisson regime.

The TDI scheme relies on the image velocity being uniform and matching the spectral rate of the TDI spectrometer. We analyzed bead velocity through the measurement region by tracking their horizontal displacement in the high-speed video data using a frame-to-frame cross correlation algorithm. These results are shown in Figure 2D. We found bead velocity averaged 1.56 mm/s with a standard deviation of 13 μm/s. We show the temporal pattern in Figure 2D because we believe it correlates with pressurization cycles in our sample pump. With perfect flow-TDI matching, signal from a 3 µm PS bead would distribute across 2-3 pixels along the TDI axis of the spectrometer (9.96x magnification produced an image width of 30 µm on the CCD, which had a pixel width of 20 µm). A histogram of the spectral widths of the 2271 events detected by our threshold algorithm, displayed in Figure 2E, shows 65% of single peak events had a spectral width of 2-3 spectra, as selected by our threshold. For the 35% of peak events larger than 2-3 spectra, we believe flow-TDI rate mismatch during the pump pressurization cycle played a role, along with contributions from beads too clustered to distinguish individual peaks. We further explored the effect of a flow-TDI rate mismatch on spectral quality and peak intensity by flowing PS beads at set pump rate while varying the TDI spectral rate, with results shown in Figure S1.

We flowed a mixed sample of PS (3 µm) and PMMA (4 µm) beads, collecting 503 event spectra at a throughput of 12 event/s (Movie S2). Figure 2F shows single event spectra from this data, with PS in cyan and PMMA in pink, as determined by their spectral profile. In the data, 17% of events consisted of mixed-bead events with peaks from both PS and PMMA. A scatterplot of peak intensity at the PMMA peak location at 2955 cm^-1^ versus the PS peak location at 3051 cm^-1^ is shown in Figure 2G. The pure-species events in this data show good separation in the scatter plot.

### Cell measurements

We demonstrated flow cytometry with fixed HeLa cells at a throughput of 32 events/s (Movie S3). To accommodate the width of HeLa cells (up to ∼40 µm), the flow cytometer detection line was configured with a 20x 0.45 NA objective, resulting in a total magnification of 4.8x to confine the cell signal to fewer pixels along the TDI axis. Figure 3A shows scattering intensity versus time for the HeLa sample under flow conditions. Figure 3B shows zoomed section of the data in 3A. Most events consisted of a single narrow peak along the temporal axis, which we attribute to the passage of one cell through the measurement region. Wider events with multiple peaks were likely due to the passage of a clumped cells. Our threshold-based algorithm selected 353 events and our peak-based algorithm selected 429 events, with an example of these selections shown by blue and orange dots, respectively, in the figure. Each event spectrum was averaged from all above-threshold event spectra for threshold-based event selection, and from the seven contiguous spectra spanning and centered on the peak position for peak-based selection. Examples of these spectra, corresponding to the lowercase alphabet index (a, b, c, …) are shown in Figure 4C. Each spectrum shows fingerprint region peaks commonly attributed to cytochromes (750 and 1585 cm^-1^), phenylalanine (1004 cm^-1^), lipids (1085 and 1458 cm^-1^), amide III bonds (1252 cm^-1^), and amide I bonds (1669 cm^-1^), among other peaks. Additionally, each event shows strong intensity in the high wavenumber region that is associated with CH_2_ vibration in lipids (2862 cm^-1^ and 2931 cm^-1^) and CH_3_ vibration in proteins (2931 cm^-1^). For these HeLa cell data and the later murine mast cell data, cytochromes have a resonant Raman response to our illumination wavelength of 532 nm, however the other molecules listed above do not. Figure 3D shows a scatterplot of maximum intensity (signal height) versus integrated intensity (total signal) for threshold-selected events. We expected the ratio of these values to distinguish single cell events from cell-cluster events in the data, and color coded this data by the number of peaks per event. The color coding showed strong grouping between event width and peak number, suggesting the ratio of peak intensity to area could be used as a gating metric to distinguish single cell events.

**Figure 3:**
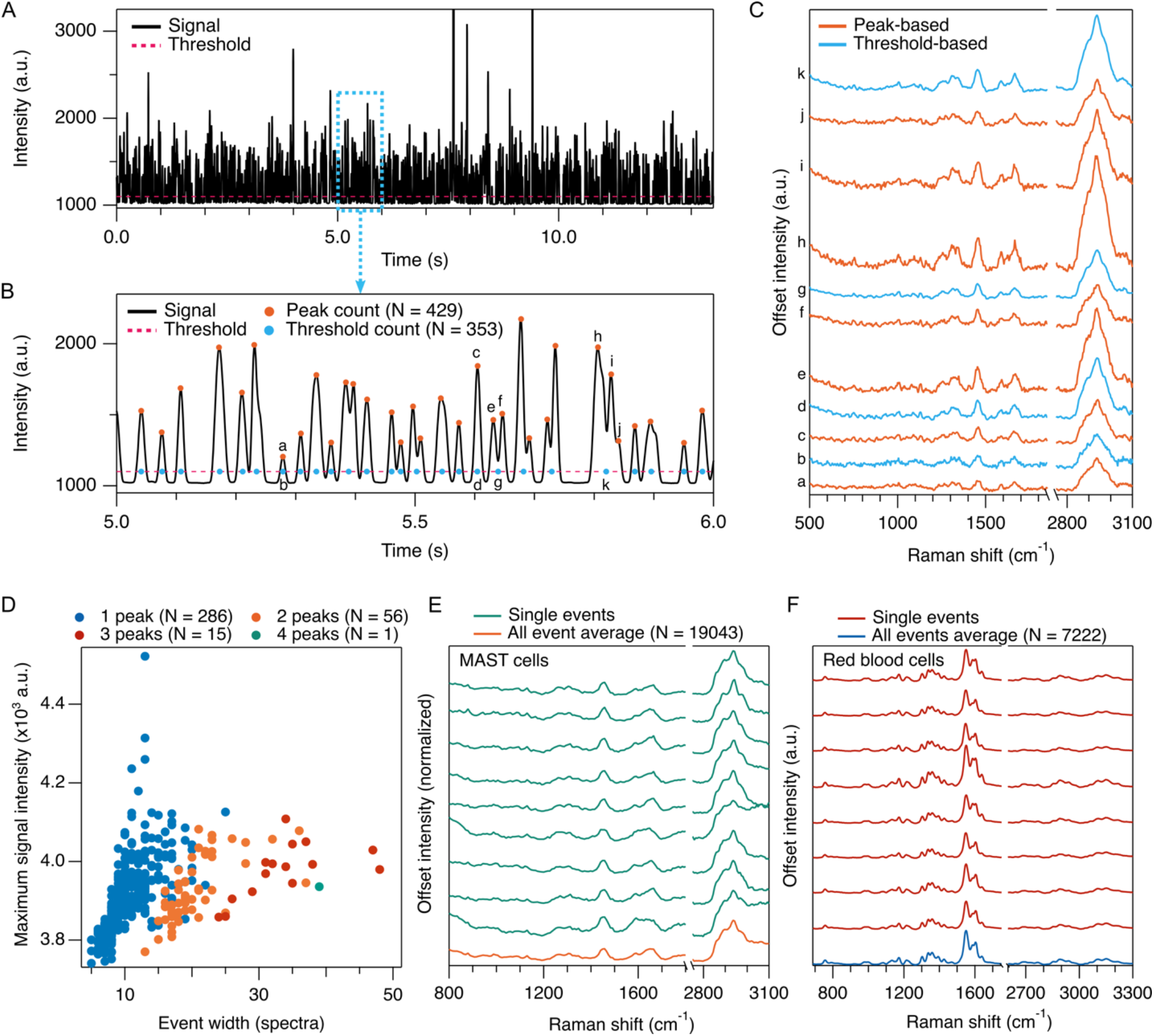
High throughput measurement of cells. **(A)** Integrated signal intensity from HeLa cells under flow. **(B)** Integrated signal intensity from 5.0 – 6.0 s showing threshold and event selection by threshold algorithm (light blue dots) and peak picking algorithm (orange dots). Throughputs measure 32 and 26 events/s, respectively. **(C)** Single event spectra for peaks indicated in B by lower case letters. **(D)** example of single cell gating using scatter plot of maximum signal intensity versus event width. Colors are assigned by intra-event peak count. **(E)** Successive single event spectra and sample average from mast cell measurement. Throughput during measurement averaged 17 events/s. **(F)** Successive single event spectra and sample average from erythrocytes. Spectral profile is dominated by the resonance Raman response of hemoglobin. Throughput during measurement averaged 26 events/s.

**Figure 4:**
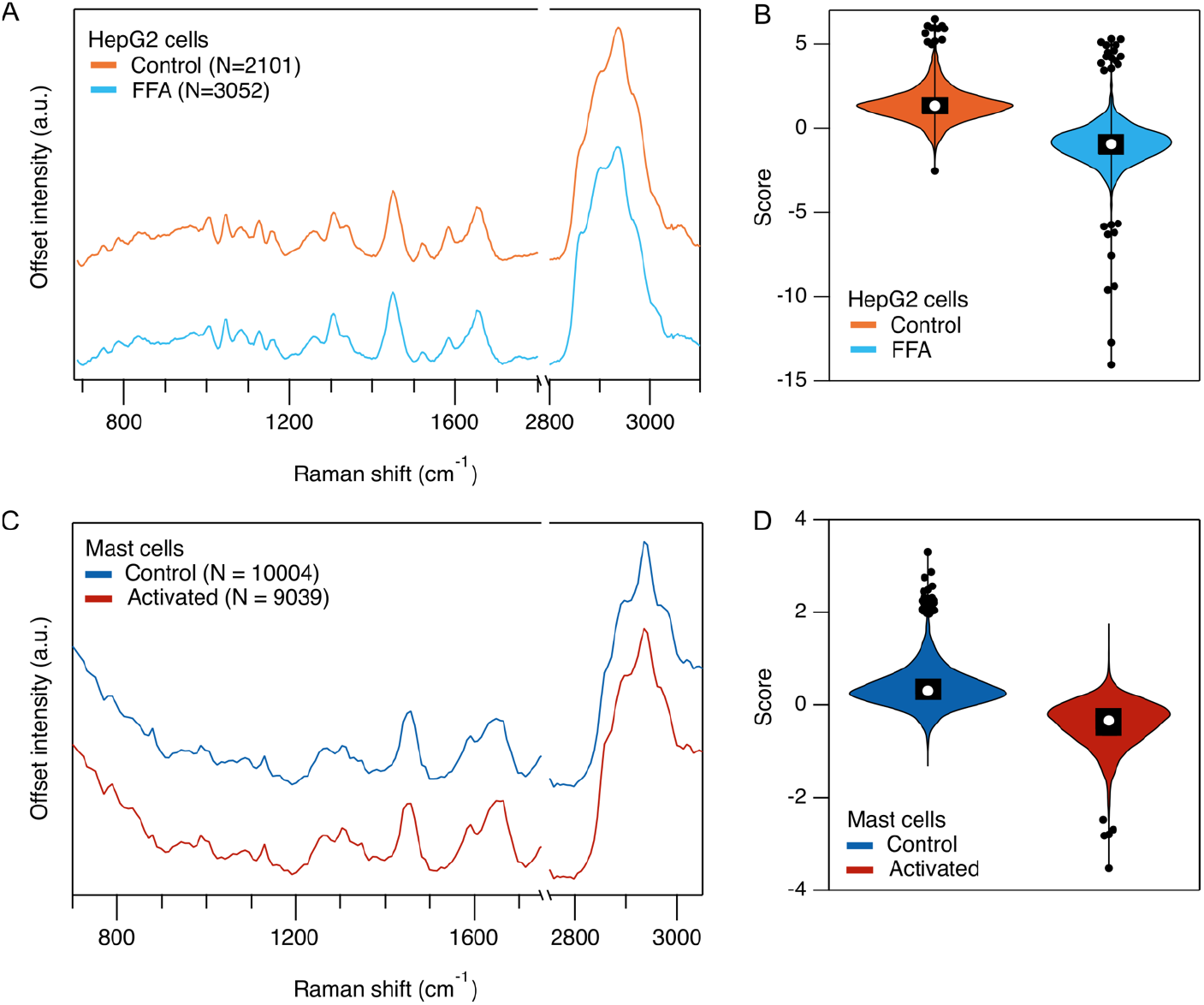
Measurement of lipid content in HepG2 cells. **(A)** Averaged spectra from HepG2 cells cultured in media with free fatty acid (FFA), vehicle control (BSA), and control conditions, as measured by our flow cytometer. **(B)** Classification of HepG2 control and FFA sample Raman flow cytometry events using linear discriminant analysis. Classification accuracy was 89%. **(C)** Averaged spectra from granulated (control) and degranulated (activated) mast cells. **(D)** Classification of mast cell Raman flow cytometry events using linear discriminant analysis. Classification accuracy was 81%. For B & D, the black boxes show interquartile range with white circles at the medians. Black circles denote far outliers.

HeLa cells are typically large, making them comparatively easy to acoustically focus, and they give a strong Raman signal. To test the robustness of the flow cytometer when measuring other cell types, we next flowed fixed murine mast cells, which typical size is between 10 and 15 µm. We have found during Raman microscopy that mast cells have a weak Raman signal and very high background autofluorescence compared to other cell types. For the mast cell measurement, we successfully captured 19043 events at 17 event/s. Example single event spectra and a sample average spectrum are shown in Figure 3E following background removal and denoising (we show the method and results of background removal and denoising in Figure S2). Finally, to demonstrate resonance Raman measurement, we flowed human fixed erythrocytes, which feature strong resonance Raman scattering from hemoglobin at our 532 nm illumination wavelength. We captured 7222 events at a throughput of 26 event/s, with single event spectra and a sample average spectrum shown in Figure 2F. With the strong resonance Raman signal in this measurement, it is likely we could have significantly increased throughput by reducing magnification power. As it was, for this acquisition, we reduced laser power to 4-6% of the level used for normal cell measurements, resulting in an intensity of 130 μW/μm^2^ at the light sheet waist. Additionally, we performed flow cytometry on live HeLa cells, with example single cell spectra shown in Figure S3.

As an application demonstration, we analyzed the lipid content of the human HepG2 hepatocyte cell line following treatment with 0.2 mM free fatty acid (FFA). Cells were cultured under two conditions. These consisted of a control sample, and a sample with FFA in the growth media using bovine serum albumin (BSA) as a carrier. More than 1800 events for each condition were analyzed by our flow cytometer, with the average spectrum from each condition shown in Figure 4A. An increased lipid signal at 2862 cm^-1^ can be observed in the FFA sample. To distinguish this fatty acid accumulation at the single event level, we applied principal component analysis-linear discriminant analysis (PCA-LDA). PCA was first applied and the first principal components (PCs) contributing 95% were selected. Then we trained a linear discrimination analysis (LDA) model on the data sets reconstructed by the selected PCs. 5/6 of the data was used as a randomly selected training set and model accuracy was validated against the remaining 1/6 of data, yielding a classification accuracy of 89%. We show the plot of LDA scores in Figure 4B. We next used our flow cytometer to distinguish granulated and degranulated murine mast cells. Mast cells are granulocytes which play a role in inflammatory immune system by releasing histamines from intracellular storage granules in response to pathogens, parasites, or allergens. Averaged spectra from all events for granulated and degranulated samples are shown in Figure 4C. PCA-LDA classification distinguished the samples with an accuracy of 81%. We show the LDA scores in Figure 4D.

## Discussion

We demonstrated TDI Raman flow cytometry at throughputs up to 32 events/s, with spectra that span the fingerprint and high-wavenumber regions. This is a significant improvement in throughput for Raman flow cytometers at this spectral bandwidth (500 – 3400 cm^-1^) and resolution (<5 cm^-1^). The spectra show peaks commonly assigned to Raman vibrational modes of phenylalanine (1004 cm^-1^), lipids (1450 cm^-1^, 2850 cm^-1^), amide III bonds (1252 cm^-1^), and amide I bonds (1669 cm^-1^), and proteins (2931 cm^-1^), as well as resonant Raman vibrational modes in cytochromes (750 cm^-1^). We were able to measure a range of mammalian cell types and sizes, including cells with high fluorescent background, and the label-free nature of Raman flow cytometry obviated the need for staining and allowed samples to be measured after minimal preparation. In our case preparation usually consisted of an optional cell fixation step, then resuspension for adherent cells, followed by a simple concentration or dilution step to adjust the suspension to an appropriate number density, which was on the order of 10^6^ cells/mL.

In our current setup, the detection speed is limited by the readout speed of the CCD camera. For the high-throughput cell and bead data presented in Figures 2 & 3, we ran our TDI CCD at its maximum spectral rate of 729 spectra/s. Sample flow speed, and thereby throughput, improves linearly with this rate, and we expect to be able to demonstrate yet higher throughputs by switching to a faster CCD, with newer commercially available models providing a 2x to 8x improvement at the time of this writing, suggesting sustained throughputs of 100 events/s and higher are possible.

Regarding flow-TDI rate matching, our data in Figure 2D & E show that our flow cytometer undergoes periodic shifts in flow rate due to the pump’s pressurization cycle. A desync between flow rate and TDI rate has two effects. First, it distributes the event signal across a greater number of pixels. As each additional pixel contributes readout noise to the event measurement, this decreases SNR. Additionally, cells that are temporally close to each other can have their signals mixed as a mismatched TDI rate will shift an accumulating pixel from an initial cell onto its temporal neighbor, increasing the rate of multi-cell events. Thus, a pump that minimizes pressure fluctuations would improve performance of the flow cytometer for both SNR and single cell event yield. A second solution could be to monitor flow speed in real time via our high-speed imaging camera and adjust TDI rate to match, though we have not examined the availability of CCDs with adjustable readout rates during acquisition.

Another improvement to our flow cytometer can be realized in its data handling. Currently we capture up to 200000 successive spectra (∼4.5 minutes) before saving data, during which data acquisition is halted. This means that large scale acquisitions such as our mast cell data are stitched together from several successive data acquisitions for analysis in post processing. Developing seamless data transfer from the TDI spectrometer to the control software would allow longer measurements. Additionally, it would allow real-time analysis and control necessary to implement Raman-activated cell sorting. We report our event throughputs as the average event rate during active measurement for an entire dataset, and did not factor in the downtime between repeats for large samples. While best practice dictates throughput should also consider such downtimes^23^, we believe focusing on throughput during measurement is a better way to judge the potential of TDI at this proof-of-concept stage. Additionally, an agitation mechanism to keep particles and cells suspended for longer measurements will be needed.

Overcoming the random inter-bead spacing described by the Poisson distribution offers a way to further increase throughput. Along similar lines, work to create uniform cell distributions along microfluidic channels has been undertaken in the field of droplet encapsulation for cells^24^, where it has been demonstrated that more uniform event distributions are possible through careful modification of the flow channel. We note that encapsulating single cells in a uniformly spaced droplet array very much resembles the encapsulation of single cell images within uniformly spaced TDI pixels, and we believe the techniques to overcome the Poisson distribution utilized in droplet microfluidics can be extended to TDI flow cytometry (with or without droplets) to further increase single cell throughput.

Though we demonstrated measurement of live HeLa cells, we did not typify cell viability following measurement. We leave the exploration of live cell yield, an important consideration for the extension of TDI to cell sorting, to future works. An inverse correlation between a cell’s viability and its optical absorption coefficient at the flow cytometer’s illumination wavelength is expected. We calculate illumination intensity in our measurements to be 0.52 – 6.4 mW/µm^2^ (depending on the sample) at the light sheet’s 7 µm beam waist for an exposure of up to 1s per cell volume. We note our previous demonstrations of live-cell compatible line-illumination Raman microscopy typically used an intensity of 1.0 – 3.0 mW/µm^2^ and a total exposure time of 2s to 5s per cell volume, which is a similar order of total exposure to our flow cytometer.

We expect the broad spectral range of TDI Raman detection to also be compatible with labelled flow cytometry using Raman tags and isotopic labels. Significant efforts have gone towards developing such tags,^25–27^ and their narrow Raman peak structures allow dozens of “colors” to be distinguished, enabling highly multiplexed labeling of intracellular content.

## Conclusion

We have demonstrated TDI-Raman flow cytometry at >30 events/s, which is a 6x improvement in throughput for cell measurement spanning the fingerprint and high wavenumber region. Our result indicates that further significant improvements to throughput can be achieved using the TDI method with modifications to camera rate and microfluidic channel design, pushing throughput above 100 cells/s without sacrificing spectral range or analyte sensitivity. We expect this to further open Raman flow cytometry to high-throughput cell analysis and cell sorting applications.

## Materials and Methods

### Cytometer design

The optical setup configuration is as described in Figure1. A 532 nm CW laser (Millenia-eV 25W, Spectra Physics) was used for the excitation light source. The beam was collimated by a beam expander that consists of two lenses (AC254-050-A-ML and AC254-200-A-ML, Thorlabs). We controlled beam power by the combination of a half wave plate (WPH10M-532, Thorlabs) and a Glan-laser calcite polarizer (10GL08AR.14, Newport). The beam size in the z-direction was individually adjusted by a pair of cylindrical lenses (f=200 and ACY254-050-A, Thorlabs) to tune the effective excitation NA, and shaped into a sheet by the combination of a cylindrical lens (LJ1567RM, Thorlabs) and a spherical lens (AC254-030-A-ML, Thorlabs). The laser passing through the quartz capillary (Noel) was collected by a lens pair (AC254-030-A-ML and AC254-075-ML, Thorlabs) having a quarter wave plate (WPQ10M-532, Thorlabs) in between. The laser beam was reflected by a mirror back to the capillary to balance the optical force on samples and align the flow path. The back-propagating beam was reflected by the polarizer to the other beam dump to prevent incidence at the laser head. The Raman scattering was collected by an objective lens (Plan Fluor ELWD 40x 0.6NA or 20x 0.45 NA, Nikon) and imaged by a tube lens (G063204000, Linos). After reflection by a 546 nm short-pass edge filter (FF01-546/SP-25, Semrock), the signal transited a relay system (ACT508-250-A-ML, Thorlabs and AC508-075-A-ML, Thorlabs) with an edge long-pass edge filter (LP03-532-RU-50, Semrock) between the relay lenses and was imaged on a spectrophotometer (CLP-100, Bunkoukeiki). After dispersion by a grating (1200 grooves/mm grating blazed at 500 nm) inside the spectrometer, the Raman spectra were image on a cooled CCD camera (PIXIS 400BR, Teledyne). High speed video imaging utilized a Memrecam Q2M (nac Imaging Technology) containing a complementary metal oxide semiconductor image sensor (ST-865, nac Image Technology).

### Flow system

We used a pressure pump (P-pump, Dolomite Microfluidics) with an integrated flow sensor for consistent sample flow. A square quartz capillary (Square Capillary 1, Nakahara Opto-Electronic Laboratory, Inc) was used at the measurement region due to the reduced background of quartz compared to glass for Raman measurement. Flow out after the capillary was routed to a waste container. Cells and particles in flow were acoustically focused by a piezoelectric transducer (Fuji Ceramics, 3.089Z6D-SY) affixed to the capillary with super glue. We drove the transducer with a 3.096 MHz sine wave output at 1 Vpp from a signal generator (WF1976, NF Corporation) and then amplified by a bipolar amplifier (HSA4101, NF Corporation).

### Data acquisition

Up to 200000 spectra were recorded per measurement, equivalent to 4 minutes and 34 seconds of continuous data acquisition, after which acquisition was halted for a few minutes as data was saved on the control PC. To acquire larger datasets, we performed multiple successive measurements.

### High-speed video analysis

We calculated bead velocity using high-speed video data that consisted of successive frames of 1920 pixel x 40 pixel images recorded at 100 fps, where the beads travelled along the 1920 pixel axis. The algorithm first flattened the images to a 1920 × 1 pixel intensity profile using a maximum value function. Using these profiles, bead velocity at frame *i* was then calculated by a cross correlation between frame *i*-5 and *i*+5, where the maximum value of the cross correlation gave the displacement in pixels during a 10-frame time interval. This displacement in pixels was then converted to distance using the camera pixel size and magnification.

### Spectral data processing

Our post-processing of spectral data consisted of four steps: pixel-to-wavenumber calibration, quartz and water baseline removal, fluorescence background removal, and noise removal. The spectrometer camera captured 1340 pixels along its spectral axis as raw data, of which 800 pixels were used to span the fingerprint, cell silent, and high wavenumber Raman regions (205 – 3320 cm^-1^ for our HeLa data), as shown in Figure S2A. Raman scattering from the quartz capillary produced small background peaks below 1000 cm^-1^, while Raman scattering from water produced a peak at 1600 cm^-1^ and a very strong and broad peak at ∼3300 cm^-1^. To isolate this baseline signal, we selected 200 spectra from event-free regions then averaged these spectra and smoothed the result with an 11-pixel binomial filter. Figure S2B shows examples of an event-average spectrum and a non-event average spectrum taken from our data shown in Figure S2A. We then subtracted this smoothed background from the raw data to remove the quartz and water Raman baseline, with the result shown in Figure S2C in light blue, which captures the average of all spectra as a line and the standard deviation from all spectra as a shaded region. Autofluorescence background from the cells shows as a parabola-like intensity across the spectral region and is a major contributor to the large standard deviation. We then used a polynomial baseline removal algorithm to remove fluorescence background from the event data, the average of which is shown as the pink spectrum in Figure S2C, with the standard deviation shown with pink shading. Next, we denoised the spectra using singular value decomposition (SVD). An example of this denoising result is shown for an example single event in Figure S2D, where the denoised spectrum was constructed from by keeping the first 52 singular values and setting the remainder to 0. The SVD scores and singular value matrix data are shown in Figures S2E and F. For experiments comparing multiple samples, the SVD was applied to a single data matrix concatenated from all samples. Pixel to wavenumber calibration was performed by measuring neat ethanol flowing in the flow cytometer and fitting the pixel values of spectral peak locations to known ethanol Raman modes at 884, 1052, 1454, and 2931 cm^-1^ with a third order polynomial. Given the weak signals in the mast cell data, we performed a 2-pixel binning along the spectral axis before post processing. This improved SNR but reduced spectral resolution to <10 cm^-1^.

### Illumination intensities

Illumination intensities were 6.4 mW/µm^2^ for mast and HepG2 cells, 4.6 mW/µm^2^ for HeLa cells, 1.3 mW/µm^2^ for microplastic particles, and 0.26 mW/µm^2^ for red blood cells. These intensities are given as the sum of the incident and retroreflected beams.

### Particle preparation

Standard microplastic concentrations were created by an initial dilution of commercial bead samples of known concentration (2.5% w/v 3 µm PS beads, Polysciences; 10% w/v 4 µm PMMA beads, Microparticles GmbH), followed by hand counting using a Neubauer chamber and an adjustment dilution as necessary. Our pure PS sample was prepared at 6 × 10^6^ beads/mL and our mixed-bead sample was prepared with equal counts PS and PMMA at a combined 1 × 10^6^ beads/mL.

### Cell preparation

HeLa cells were cultured on 25 mm^2^ cell T-flask in Dulbecco’s Modified Eagle Medium (DMEM, 043-30085, Wako-Fujifilm) supplied with 10% fetal bovine serum (FBS, S1780-500U, Biowest), and 1% penicillin-streptomycin-glutamine solution (15140122, Gibco) in a CO_2_ incubator at 37 °C.

HepG2 cells were cultured in minimum essential medium (MEM, 51200038, Thermo fisher Scientific) supplemented with 10% fetal bovine serum (FBS, S1780-500U, Biowest), and 1% penicillin-streptomycin-glutamine solution (15140122, Gibco) for in the CO_2_ incubator at 37 °C. On day two and three, we added different concentrations of FFA solutions (0 or 0.2 mM) to the HepG2 cell cultures. The FFA solutions were formulated by dissolving palmitic acid (P9767, Sigma-Aldrich) and oleic acid (O7501, Sigma-Aldrich) in a mole ratio of 1:2 into culture medium containing 0.2 mM FFAs-free BSA. For measurement, HepG2 cells under each condition were collected and into a 4 mL jar and the medium was replaced with live cell imaging solution (A14291DJ Thermofisher).

### Culturing of BMMC’s and degranulation

Bone marrow-derived mast cells (BMMCs) were generated by culturing bone marrow cells derived from a 10-week-old female C57BL/6NCrSlc mouse (Japan SLC) in RPMI-1640 medium (R8758-500ML, SIGMA) supplemented with 10% fetal bovine serum, 10 mM HEPES, 1 mM sodium pyruvate, 55 µM 2-mercaptoethanol, 100 U/mL penicillin, 100 µg/mL streptomycin, and 10 ng/mL recombinant mouse IL-3 (554579, BD Pharmingen) for 4 weeks.

5 × 10^6^ BMMCs were incubated with 200 ng/mL anti-TNP IgE (554118, BD Biosciences) for 2 hours. After washing with 1× PBS, the cells were resuspended in 1× Tyrode buffer and then stimulated with or without 3 ng/mL TNP-BSA (LG-1117, LSL) for 1 hour.

### Erythrocytes

Erythrocytes and human serum were obtained from the Japanese Red Cross (research ID:25J0143) with written informed consent. Erythrocytes were cultivated at 2% hematocrit in complete medium, which consists of RPMI-1640 medium containing 2.5% human serum, 2.5% AlbuMAX II (Life Technologies Carlsbad, USA), 25 mM HEPES, 0.225% sodium bicarbonate, and 0.38 mM hypoxanthine supplemented with 10 ug/ml gentamicin. RBCs were incubated under low-oxygen conditions (90% N_2_, 5% CO_2_, and 5% O_2_).

### Cell fixation

Cells were fixed in 4% paraformaldehyde and rinsed three times with phosphate buffer solution.

### Code base

Processing of spectral and image data was performed using custom code in Igor Pro (Wavemetrics) and Matlab (Mathworks).

### Statistical analysis

Interquartile range and far outliers were calculated using the Tukey method in Igor Pro (Wavemetrics).

## Acknowledgements

The authors thank Dr. Kazuki Bando and Ryutaro Ohata for helping optical setup construction.

## Funding

This work was supported by JST-Mirai Program Grant Number JPMJMI20G3, Japan (KF).

This work was also supported by JST COI-NEXT Grant Number JPMJPF2009 (KF).

This work was also supported by Japan Society for the Promotion of Science for a JSPS International Research Fellowship 22KF0235 (MLindley, KF).

## Author contributions

Project administration: KF

Conceptualization: KF

Methodology: KF, MLindley,TK

Resources: MLi, JQ, TY, SI, KM

Investigation: MLindley, TK, SD

Visualization: MLindley,TK

Supervision: KF

Writing—original draft: MLindley, TK, KF

Writing—review & editing: MLindley, TK, SD, MLi, TY, SI, KM, KF

## Competing interest statement

MLindley, TK, and KF have patent application related to this work (PCT/JP2024/003220). K.F. is a co-founder and the chief technology officer (CTO) of Millde Co., Ltd. The other authors have no competing of interests.

