## Supplementary Materials for "Raman flow cytometry using time delay integration"

bioRχiv

**Matthew Lindley,** [a,†,‡] **Toshiki Kubo,**[a,b,†] **Stéphanie Devineau,**[a,c] **Menglu Li,**[a,§] **Jing Qiao,**[a] **Takuya Yashiro,**[d,e] **Shiroh Iwanaga,**[f] **Kazuyo Moro,**[d,e,g] **and Katsumasa Fujita**[a,e,h,*]
[a]Department of Applied Physics, Graduate School of Engineering, Osaka University, 2-1 Yamadaoka, Suita, Osaka 565-0871, Japan
[b]Department of Dermatology, Course of Integrated Medicine, Graduate School of Medicine, Osaka University, 2-2 Yamadaoka, Suita, Osaka 565-0871, Japan
[c]Université Paris Cité, CNRS, Unité de Biologie Fonctionnelle et Adaptative, 75013 Paris, France
Department of Molecular Protozoology, Research Institute for Microbial Diseases, Osaka University, 3-1 Yamadaoka, Suita, Osaka 565-0871 Japan
[d]Laboratory for Innate Immune Systems, Graduate School of Medicine and IFReC, Osaka University, 2-2 Yamadaoka, Suita, Osaka 565-0871, Japan
[e]Institute for Open and Transdisciplinary Research Initiatives, Osaka University, 2-1 Yamadaoka, Suita, Osaka 565-0871, Japan
[f]Department of Molecular Protozoology, Research Institute for Microbial Diseases, Osaka University, 3-1 Yamadaoka, Suita, Osaka 565-0871 Japan
[g]Laboratory for Innate Immune Systems, RIKEN-IMS, 1-7-22 Suehiro-cho, Tsurumi-ku, Yokohama, Kanagawa 230-0045, Japan
[h]AIST-Osaka University Advanced Photonics and Biosensing Open Innovation Laboratory, National Institute of Advanced Industrial Science and Technology (AIST), 2-1 YamadaokaSuita, Osaka 565-0871, Japan

[†]These authors contributed equally to this work

### Supplementary text

#### Effects of flow rate and spectral rate matching

To achieve high SNR and temporal resolution, it was important that the velocity of the particle or cell image across the CCD face matched the TDI spectral rate. We demonstrated the effect of image velocity and TDI rate matching in Figure S1. For this demonstration, polystyrene beads suspended in water were flowed at 1.5 μL/min within the capillary. We then took data while varying the TDI rate from 435 – 505 spectra/s. Figure S1A shows this data as Raman scattering intensity from the sample at a Raman shift of 1004 cm$^{-1}$, which is a strong Raman scattering peak of polystyrene. The measurement of a single bead or bead clump appears as a peak in this temporal data. When the bead image velocity (dependent on flow rate) matched the TDI rate, the bead signal was confined to just a few pixels along the TDI axis, resulting in a narrow and intense peak. If the bead image velocity and spectral rate was mismatched, this same signal distributes across a larger number of TDI axis pixels, resulting in a motion-blurred peak that is temporally broader and weaker in maximum intensity.

Based on prior analysis of bead velocity during flow using our high-speed video, we expected a TDI rate of 465 spectra/s to produce the best data for a flow rate of 1.5 μL/min. Our data show narrow and intense peaks when the TDI rate is at both 465 and 475 spectra/s, suggesting the optimal rate may lie between these values. The data show that as the TDI rate shifts farther from the optimal value, the temporal peaks begin to broaden significantly, as expected. Raman spectra of the first four events for each flow condition are shown in Figure S1B, where the start and stop of each single event was selected by a rising and falling intensity threshold, respectively. An all event average spectrum for each sample is shown in Figure S1C. Per-event spectra were then averaged to produce a single spectrum for each event. Histograms of peak intensity at 1004 cm$^{-1}$ for all events are shown in Figure S1D. These data clearly show that spectral peak intensity maximizes as the match between spectral rate and flow rate improves.

### Supplementary Figures

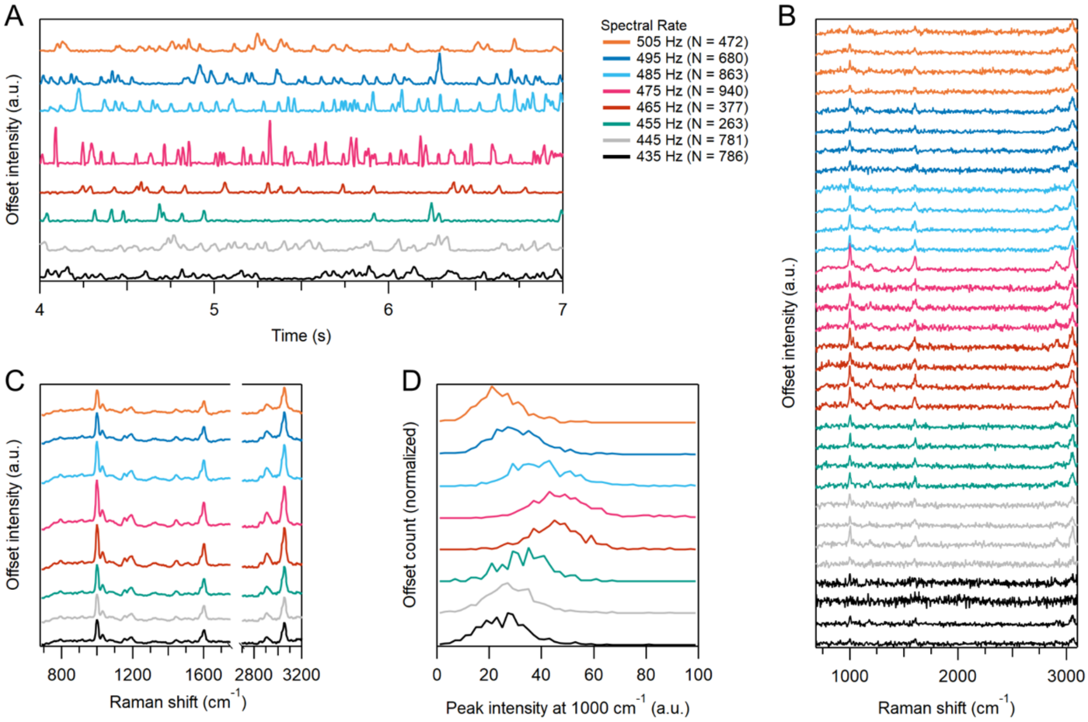


**Figure S1: Effects of flow and spectral rate matching on signal.** (A) Intensity of beads signal at 1004 $cm^{-1}$ as TDI spectral rate is varied while sample flow rate is kept constant. Dashed black lines correspond to the event selection threshold. (B) Corresponding single event spectra measured by flow cytometer, without denoising, treated with polynomial baseline removal. (C) Sample averages. (D) Peak intensity at 1004 $cm^{-1}$ vs baseline.

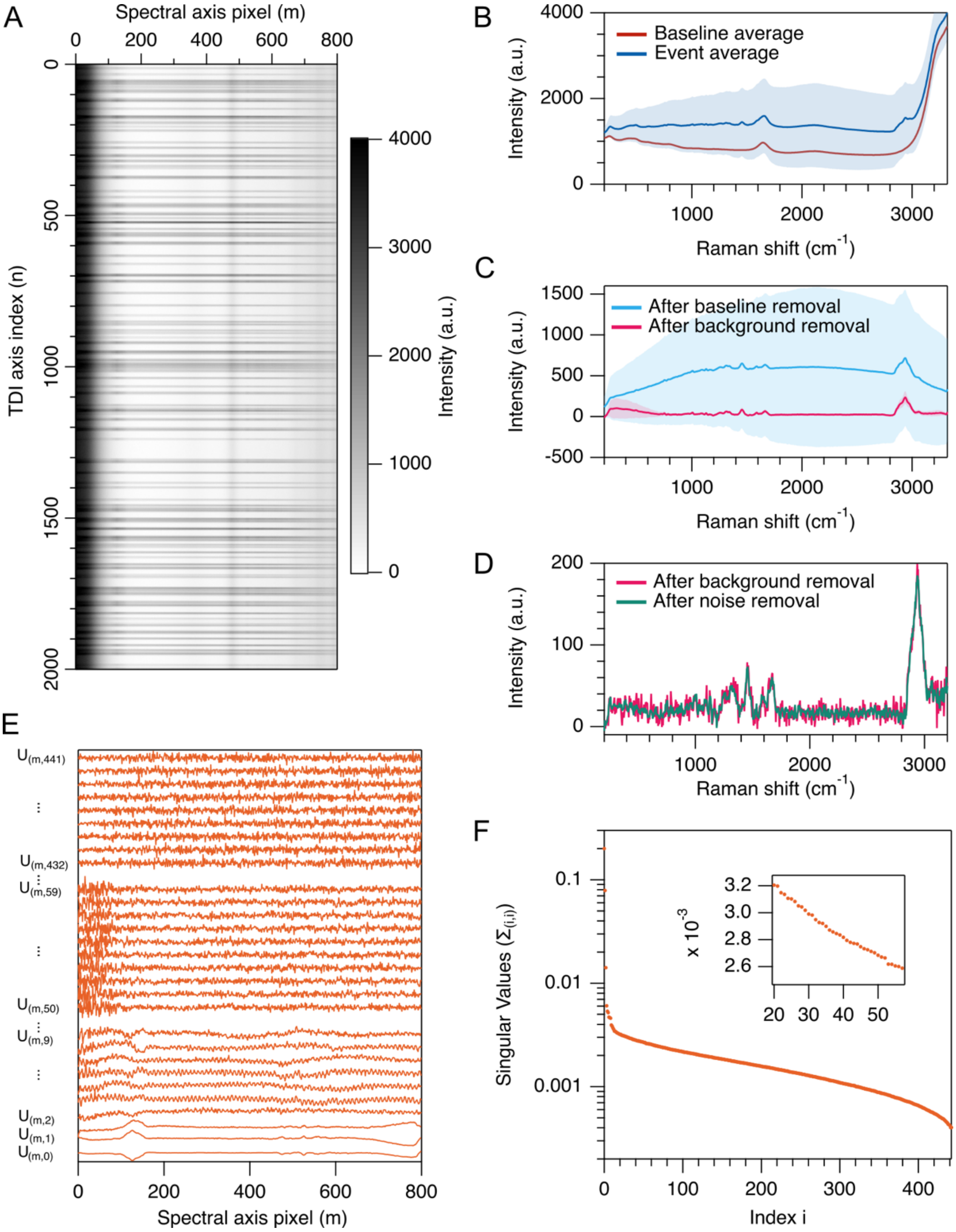


**Figure S2: Post processing for scattering baseline, fluorescence background, and noise removal.** (A) unprocessed data from measurement of HeLa cells. (B) Averages from all events and 200 baseline (non-event) spectra. Standard deviation is shown by the shading. (C) Averages after quartz and water peak removal (events – baseline) and fluorescence background removal. (D) A single event spectrum before and after SVD noise removal. (E) SVD noise removal U matrix. (F) SVD noise removal Σ matrix.

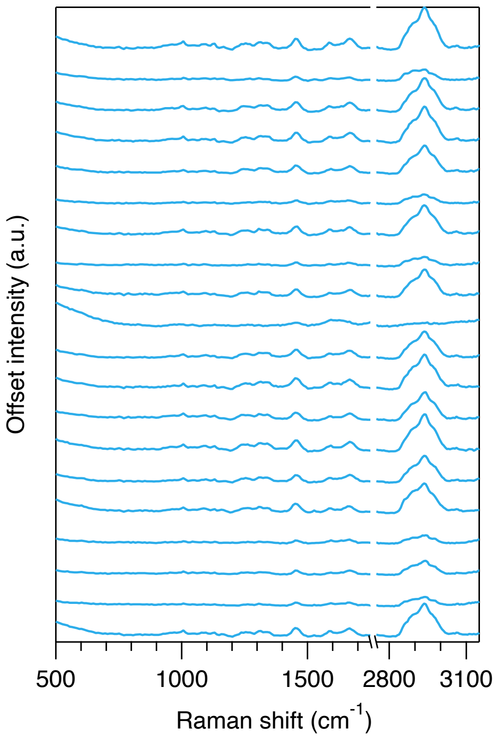


**Figure S3: Single event spectra from measurement of live HeLa cells.**

**Movie S1  Raman spectra and high-speed camera images of polystyrene (PS) beads.**

**Movie S2  Raman spectra and high-speed camera images of PS and polymethyl methacrylate (PMMA) beads mixture.**

**Movie S3 Raman spectra and high-speed camera images of fixed HeLa cells.**
